# Multiscale curvature modulates epithelial architecture and nuclear organization

**DOI:** 10.64898/2026.01.12.698973

**Authors:** Marine Luciano, Quentin Vagne, Guillaume E. Pernollet, Guillaume Salbreux, Caterina Tomba, Aurélien Roux

**Affiliations:** Department of Biochemistry, University of Geneva, Quai Ernest Ansermet 30, Geneva, CH-1211, Switzerland; Department of Genetics and Evolution, University of Geneva, Quai Ernest Ansermet 30, Geneva, CH-1211, Switzerland; CNRS, INSA Lyon, Ecole Centrale de Lyon, Universite Claude Bernard Lyon 1, CPE Lyon, INL, UMR5270, 69621 Villeurbanne, France

## Abstract

Curved geometries are a defining feature of epithelial tissues, yet how cells integrate curvature cues across spatial scales remains unclear. Here, we combined wavy hydrogels with inducible self-rolling substrates to readily and independently impose local and large-scale curvatures on epithelial monolayers, recapitulating key geometric features of bronchiolar epithelia. By controlling the orientation of local curvature relative to the large-scale curvature set by the tube axis, we show that multiscale curvature induces scale-dependent and anisotropic remodeling of cell shape, nuclear organization, and tissue thickness. In contrast, nuclei maintain a robust and conserved three-dimensional geometry, with curvature primarily regulating nuclear orientation rather than shape. This hierarchical induction of specific changes to the tissue architecture shows that multiscale curvature sensing is a fundamental physical principle governing epithelial architecture.

## Introduction

Curvature is a fundamental geometrical feature of living systems, present across all levels of biological organization, from nanometer-scale membrane deformations to organ-scale tissue folding^1,2^. While long regarded as a passive outcome of morphogenetic processes^3,4^, curvature is now recognized as an active biophysical signal that shapes cellular organization and tissue function^2^. In particular, epithelial tissues in the human body are characterized by highly curved structures spanning a wide range of spatial scales, from micrometer-scale surface undulations to organ-scale tubular and lobular architectures^5^. A striking example is found in the respiratory tract, where bronchioles and alveoli exhibit complex luminal topographies with continuous axial and radial curvature, forming highly folded epithelial layers critical for gas exchange. Such geometrical features not only guide tissue morphogenesis but also modulate physiological processes including airflow, airway resistance, and pathogenesis in conditions such as asthma or chronic obstructive pulmonary disease (COPD)^6,7^. Beyond being a geometric outcome of morphogenesis and patho-physiological processes, tissue curvature has emerged as a fundamental physical cue that shapes cell organization, force balance and collective behaviour^8^. Over the past decade, increasing evidence shows that curvature actively regulates cellular processes, including cell shape, polarity, proliferation, migration and fate specification. For example, epithelial cells respond distinctly to positive and negative curvatures. Positive geometries promote cell spreading, cytoskeletal tension, and nuclear flattening, whereas negative environments induce apical constriction and actomyosin accumulation ^9–11^. The nucleus, in particular, is highly sensitive to curvature-induced changes in cell shape and force distribution. Curved substrates have been shown to drive pronounced nuclear deformations through cytoskeletal prestress, leading to changes in nuclear height, volume and chromatin organization^12^. Epithelial cells cultured on wavy substrates actively sense curvature and undergo pronounced nuclear mechanoadaptation^13^. These curvature-dependent nuclear adaptations are accompanied by modifications in lamin composition and spatial patterns of cell proliferation, highlighting the nucleus as a central mechanosensitive organelle linking geometry to epithelial function^9,12,14^.

To replicate such curved environments in vitro, a variety of biomimetic systems, including patterned hydrogels^15–18^ or curved scaffolds like bowl-shaped microwells^10,19^, have been developed to probe curvature sensing at single-cells and tissue scale. These approaches revealed that epithelial cells can generate supracellular actomyosin structures in response to curvature^17,20^, driving coordinated nuclear elongation.

Beyond static geometries, recent studies have revealed that dynamically changing curvature elicits active nuclear protection mechanisms involving actin and vimentin filaments underscoring the ability of epithelial cells to adapt nuclear architecture to fluctuating mechanical landscapes^21^. Self-rolling has enabled the rapid generation of tubular epithelial geometries^22,23^, providing access to dynamic curvature induction and revealing transient mechano-osmotic responses to anisotropic folding^24^.

Importantly, curvature is increasingly linked to pathological processes^25^. In epithelial tissues, curvature gradients have been correlated with oncogenic transformation. For instance, three-dimensional (3D) imaging and modeling of pancreatic ducts demonstrated that duct curvature influences the directionality of epithelial growth, with small ducts favouring exophytic growth, and larger ducts undergoing endophytic deformation^26^. These findings suggest that tissue curvature should be considered as a key physical factor contributing in epithelial tumorigenesis.

In vivo, epithelial tissues are rarely exposed to a single curvature scale. Instead, they experience complex geometries in which local surface undulations coexist with large-scale tissue curvature, often with pronounced anisotropy. Whether epithelial tissues integrate curvature cues independently at each scale or combine them through a hierarchical, multiscale mechanism remains unknown. Addressing this question has been limited by the lack of experimental systems allowing controlled and simultaneous manipulation of curvature across spatial scales in soft, biologically relevant environments compatible with live epithelial tissues. Here, we combined wavy polyacrylamide hydrogel with inducible self-rolling polydimethylsiloxane (PDMS) multilayers to generate tubular epithelial monolayers with local folding. These geometries recapitulate key features of curved epithelial tissues, such as bronchial tubes, and are compatible with live cell imaging and morphometric analyses. Importantly, the orientation of local curvature relative to the tube axis can be precisely controlled, enabling systematic exploration of curvature anisotropy, while dynamic curvature configurations can be directly compared to static conditions on wavy two-dimensional hydrogels.

## Results

### Engineering epithelial tissues with controlled multiscale curvature via self-rolling

The formation of tubular structures whose inner lumen features a wavy surface was generated as follows: large-scale curvature was induced by triggering the self-rolling of a pre- strained PDMS bilayer^23,24^. This PDMS bilayer was formed by spin-coating successively two layers of PDMS, the second one containing 48% of oil. The oil was subsequently extracted using toluene, creating an internal strain within the PDMS bilayer. Importantly, the PDMS bilayer was assembled on a gelatin-coated region, which prevented PDMS adhesion in central areas while maintaining strong peripheral attachment to the underlying PDMS. This configuration facilitated localized release and self-rolling upon cutting.

Local curvature was generated by photopolymerizing polyacrylamide hydrogels on top of the PDMS bilayer using a photoiniator and a striped optical photomask^9^, creating wavy hydrogels with a wavelength of 100 µm **(Fig. 1A-C)**. To ensure robust adhesion between the hydrogel and PDMS layers, a benzophenone-based surface activation strategy was employed, enabling covalent bonding between PAAm and PDMS substrates^27^. This strong interface withstands substantial mechanical deformation during the rolling process.

**Figure 1.**
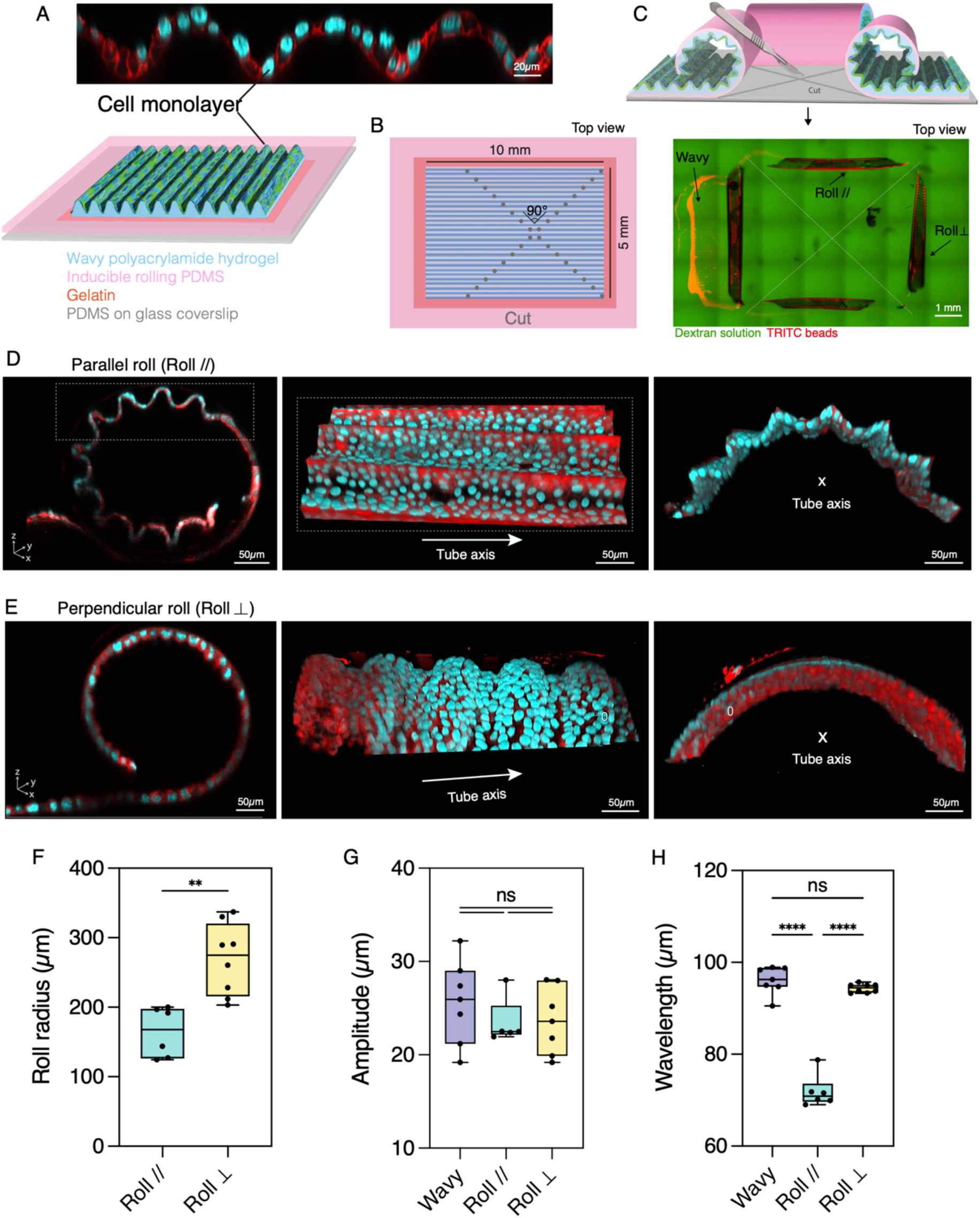
Engineering epithelial monolayers under controlled multiscale and anisotropic curvature. **(A)** Bottom – Schematic of the multi-layer substrate composed of glass slide with PDMS (gray); inducible PDMS sheets (pink); central gelatin patch (orange), wavy polyacrylamide hydrogel (light blue), cell monolayer (green). Top – Orthogonal view of a MDCK epithelial cells on wavy hydrogels. Cell membrane (deep-red CellMask, red) and nuclei (Hoechst, cyan). **(B)** Schematic of the system top view before cutting. Gray dotted lines represent the cut axis. **(C)** Top: Schematic of the self-rolling of the substrate upon cutting leading to have on the same substrate : two parallel rolls, two perpendicular rolls and one wavy zone. Bottom: Top view of the subtrate upon cutting without cells. Dextran solution (green) and TRITC beads inside the hydrogel (red). **(D)** Confocal images of a parallel roll with cells. Cell membrane (deep-red CellMask, red), and nuclei (Hoechst, cyan). **(E)** Confocal images of a perpendicular roll with cells. Cell membrane (deep-red CellMask, green), and nuclei (Hoechst, cyan). Distribution of roll radius **(F)**, amplitude **(G)** and the wavelength **(H)** with cells after cutting for wavy (light purple), parallel (cyan) and perpendicular (yellow) configurations. The roll radius is taken at the mid distance between crest and valley. **p<0.01 and ****p<0.0001.

Wavy PAAm hydrogels were first functionalized with human fibronectin (FN) and Madin-Darby Canine Kidney (MDCK) cells were seeded at confluency. After 48 hours in culture, the PDMS bilayer, along with the attached hydrogel and epithelial monolayer, was incised to release the pre-strain and initiate the rolling process **(Fig. 1B-C)**. This led to the spontaneous formation of tubular structures with a wavy inner surface, enabling controlled investigation of epithelial responses to abrupt curvature transitions across multiple spatial scales. To explore curvature anisotropy, the incision was performed in a crosswise manner, generating tubes with undulations aligned with **(Fig. 1D)** or perpendicular **(Fig. 1E)** to the tube axis, respectively named parallel rolls and perpendicular rolls. The unrolled configuration is named wavy in the following. The resulting tube radii were reproducible and fell within the range of curvature observed in native epithelial ducts, including bronchial and glandular structures ^5^.

Quantitative geometric analysis revealed that tube radius depended on the orientation of undulation pattern, with smaller radii for parallel rolls r = 164 ± 36 µm compared to the perpendicular rolls r = 269 ± 52 µm **(Fig. 1F)**, probably reflecting hydrogel’s anisotropic resistance to rolling. This is supported by the fact that in experiments without hydrogels ^24^, the tube radii were even smaller than that. In contrast, the amplitude of the wavy pattern remained largely unchanged upon rolling **(Fig. 1G)**, whereas its effective wavelength λ was selectively altered in an orientation-dependent manner. Indeed, we obtained λ = 96 ± 3 µm for wavy substrates, λ = 72 ± 4 µm for parallel rolls, and λ = 94 ± 1 µm for perpendicular rolls **(Fig. 1H)**.

Together, these results establish a robust experimental platform that enables controlled and reproducible exposure of epithelial tissues to multiscale and anisotropic curvature. Consistent with previous observations on homogeneous self-rolling substrates, curvature induction is accompanied by lateral compression during rolling, estimated to reach ∼15% ^24^. In the presence of a wavy hydrogel layer, this compression is redistributed in an orientation-dependent manner, resulting in distinct tube radii and wavelengths in the parallel and perpendicular configurations.

### A global geometric framework to describe multiscale curvature

To quantitatively characterize how local and large-scale curvature combine upon rolling, we first developed a method to mathematically describe the complex three-dimensional (3D) shapes of the different substrates after curvature induction **(Supplementary Fig. 2)**. We focused on quantifying the principal directions of curvature from confocal reconstructions: they revealed complex surface geometries resulting from the interweaving of large-scale cylindrical curvature and local undulations **(Fig. 2A)**. We reconstructed 3D meshes from the segmented basal (i.e. the cell-substrate interface) and apical surfaces of each substrate type **(Fig. 2A and Supplementary Fig. 2)**. Representative meshes of the wavy, parallel, and perpendicular geometries are shown in Fig. 2A. While curvature tensors could be computed locally from the raw mesh data, it often yields noisy results due to mesh irregularities. To obtain well resolved curvature profiles, we adopted a global surface-fitting strategy using idealized mathematical models that capture the essential geometric features of each substrate **(Fig. 2B, and Supplementary Section 1)**. A custom analysis pipeline was developed to fit this idealized mathematical surface to the segmented meshes for each substrate **(Fig. 2D and Supplementary Section 2)** and global shape parameters were extracted for all substrate types **(Fig. 1F-H)**. This approach enabled robust extraction of local and large-scale geometric parameters, including effective wavelength (𝜆) and amplitude (A), while minimizing noise inherent to local curvature tensor estimation **(Fig. 2C)**. For rolled substrates with large-scale curvature, we additionally identified the radius R (with 1/R the large-scale cylindrical curvature), and the wave orientation angle (𝜙), with 𝜙 = 0 for parallel and 𝜙 = 90° for perpendicular alignment relative to the tube axis.

**Figure 2.**
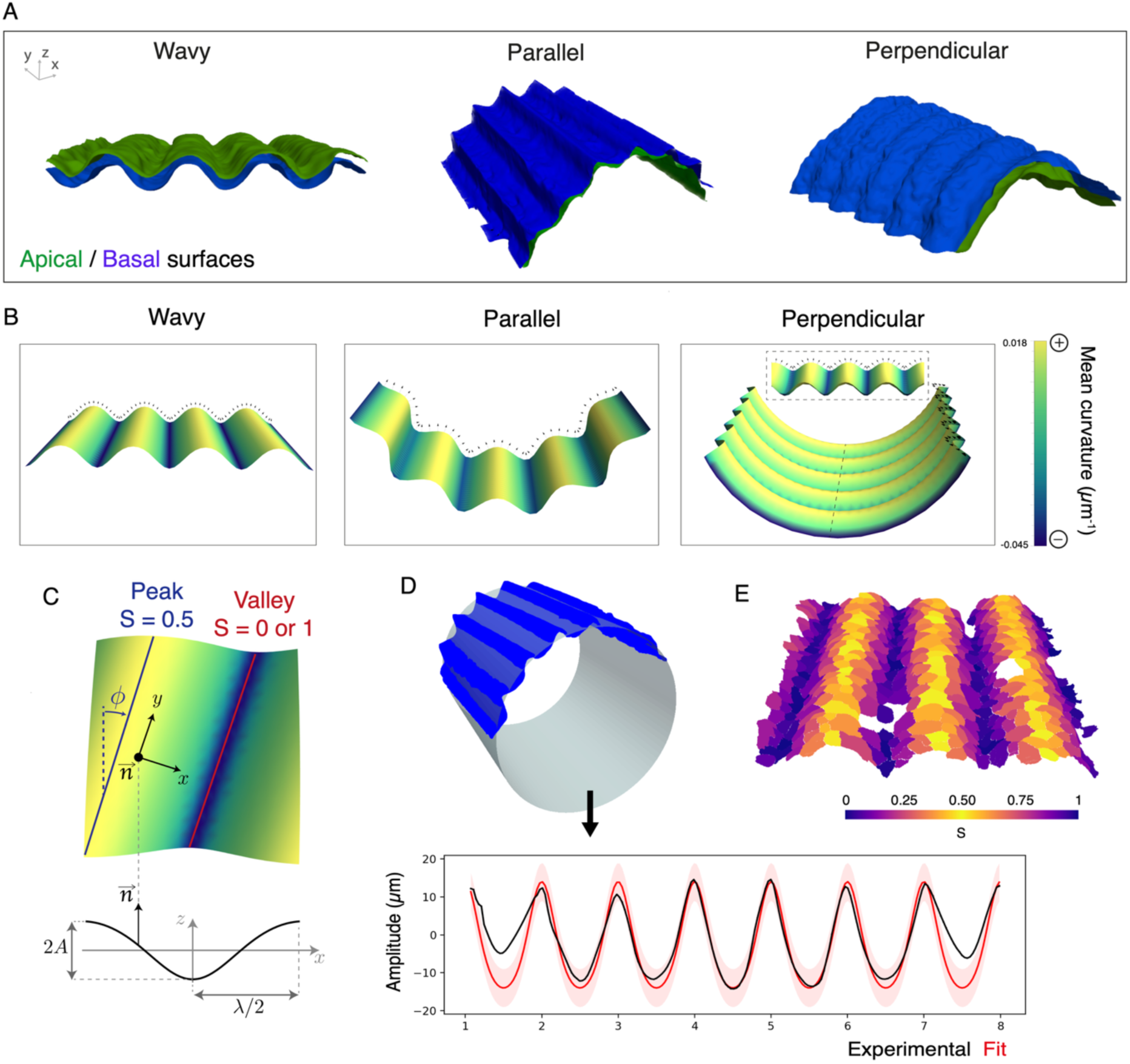
Quantitative geometric framework for multiscale substrate curvature analysis. **(A)** Examples of segmented apical (green) and basal (blue) meshes in representative wavy and rolled (parallel, perpendicular) configurations. **(B)** Idealized mathematical surfaces used to fit the substrate shape, in typical wavy and rolled (parallel, perpendicular) configurations. The surfaces are colored according to the local, analytically computed, mean curvature. Black arrows represent the local normal vector. **(C)** Illustration of the main geometrical parameters characterizing the substrates. (Top) The tilt angle 𝜙 (for cylindrical substrates) of the oscillations, the local X, Y, Z directions aligned with the average plane/cylinder and the main directions of curvature. (Bottom) Oscillation profile described by its amplitude A and its wavelength 𝜆. **(D)** Fitting procedure, here in the rolled configuration. A cylinder is fitted to the basal mesh (blue), which is then used to unwrap the oscillation profile and collapse it onto one dimensional representation showing the average amplitude (black line) as a function of the (non-periodic) coordinate S. The idealized mathematical profile (red line, +/- 5𝜇m tolerance shown in transparent red) is fitted to the experimental one. **(E)** Typical wavy substrate after the fitting procedure is completed. Segmented apical sides of cells are shown. Each cell is colored according to its (periodic) S coordinate.

Furthermore, the continuous nature of the fitted surfaces allows to extract a precise curvature tensor at every point of the idealized surface, which contains the information of the two main directions of curvatures, with their associated main curvature values 𝜅_1_ and 𝜅_2_. From these, we computed the mean curvature 𝜅 = (𝜅_1_ + 𝜅_2_)/2, which quantifies the local mean curvature **(Supplementary Fig. 1)**.

Fitted surfaces also allowed to extract local reference vectors: a normal vector 𝑛/⃗ which is orthogonal to the large-scale substrate shape, unperturbed by the wavy pattern, and two directions orthogonal to 𝑛/⃗, X and Y **(Fig. 2C and Supplementary Section 1)**. X is the direction going from valleys to valleys, while Y is in the direction of valleys/ridge lines. We defined a periodic coordinate S along direction X **(Fig. 2E)**, to represent the position relative to peaks and valleys. At valleys, S=0 or 1, while at peaks, S=0.5.

Importantly, this formal geometric description establishes a unified framework to compare substrates in which local and large-scale curvature are combined, and to analyze how orientation-dependent rolling constraints reshape the effective geometry experienced by epithelial tissues. Together, this analysis provides a quantitative geometric basis for investigating how epithelial cells respond to multiscale curvature, and sets the stage for linking emergent geometry to nuclear and tissue-scale organization.

### Cellular morphology reflects integration of multiscale curvature cues

We next asked how epithelial cells interpret this complex geometry at the cellular level. In particular, we investigated whether cellular morphology and positioning reflect the multiscale curvature constraints imposed by the underlying geometry.

To determine how epithelial cells respond to induced multiscale curvatures, we first extracted apical cell contours from confocal images using a standardized segmentation workflow **(Fig. 3A)**, enabling measurement of cell apical area **(Supplementary Fig. 2)**, tissue thickness, apical surface orientation, and width along principal axis **(Supplementary Section 3)**. Apical cell area varied markedly with curvature at small length-scales, in all three conditions (wavy, parallel and perpendicular): cells were smaller in the valleys (S = 0 or 1) and larger on the crests (S = 0.5) **(Fig. 3B)**. Also, a significant reduction of the average apical area was observed upon rolling **(Fig. 3C)**.

**Figure 3.**
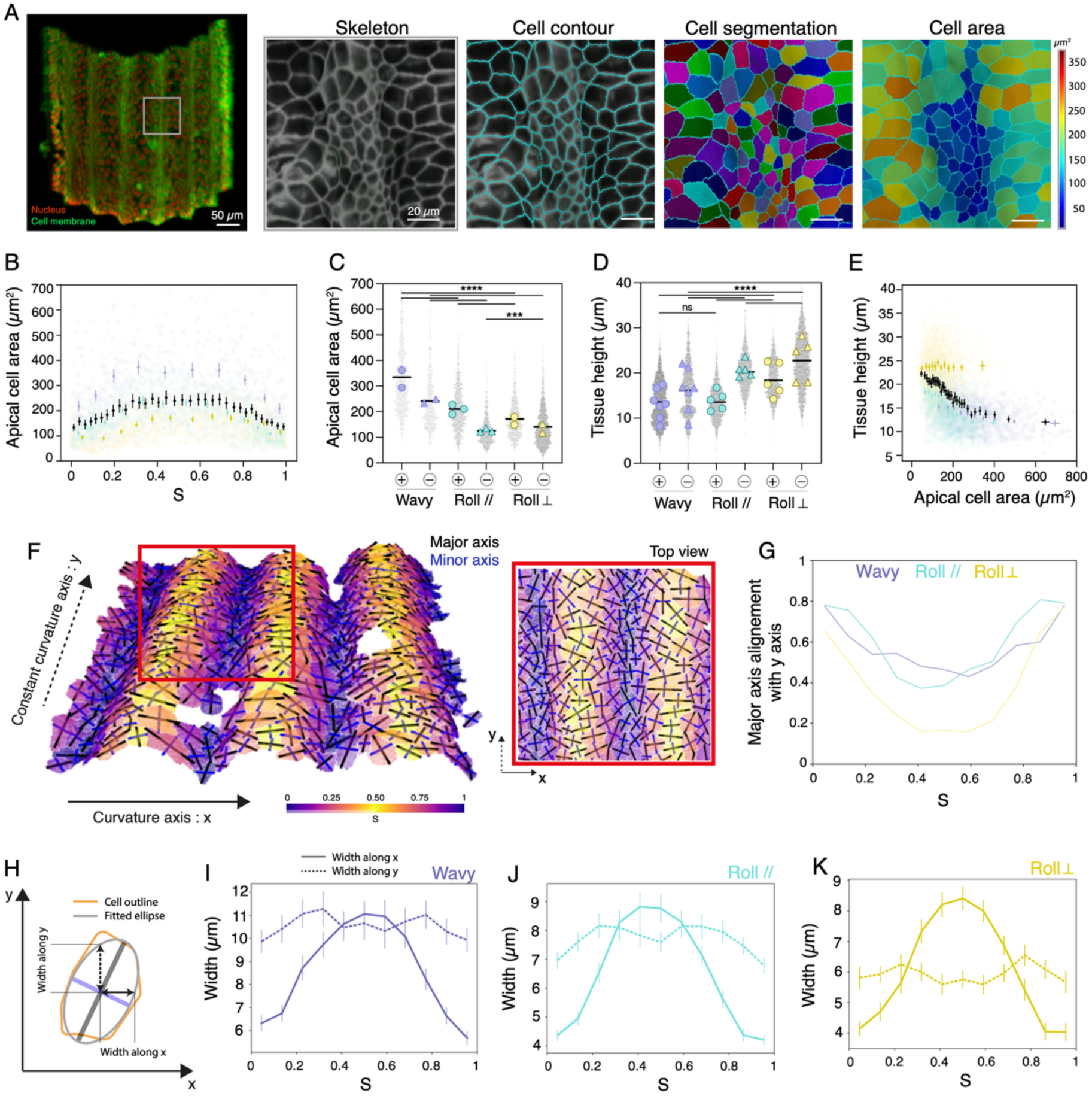
Curvature-dependent regulation of epithelial cell shape and orientation. **(A)** Left – Representative confocal image of a half parallel roll. Cell membrane (deep-red CellMask, green) and nuclei (Hoechst, red). Zoom – typical image of the projection of the cell membrane signal giving the skeleton; cell contour; cell segmentation and color-coded cell area. Quantification of the apical cell area **(B)** in function of the parameter S and **(C)** for negative and positive zone across wavy, parallel and perpendicular tissues. **(D)** Quantification of the tissue height in function of the parameter S. **(E)** Relationship between tissue height and apical cell area. **(F)** Typical 3D wavy substrate with segmented cells showing the curvature axis (X axis, dotted arrow) and the constant curvature axis (Y axis, full arrow). Black lines represent cell major axis and blue lines represent cell minor axis. Each cell is colored according to its (periodic) S coordinate. **(G)** Distribution of major axis alignment with the constant curvature axis, Y for wavy and rolled structures. **(H)** Schematic representation showing the cell width along Y and X. Cell width along Y and X direction in function of S for **(I)** wavy, **(J)** parallel and **(K)** perpendicular tissues. ***p<0.001 and ****p<0.0001.

We next measured the tissue thickness, defined as the vertical distance measured between the segmented apical and basal surfaces and along the normal vector 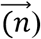 defined above. Across all conditions, our results showed that tissue thickness was consistently larger in negative curvature (valleys) and reduced on positives curvature (crests) **(Fig. 3D)**, in good agreement with previous observations on static wavy surfaces^9^. Notably, tissue thickness was globally increased in rolled configurations compared with wavy controls **(Fig. 3D)**, in good agreement with the observation of a transient swelling of cells upon induction of rolling^24^. However, tissue thickness exhibited a behavior opposite to that of projected cell area, with an overall global anticorrelation between the two parameters **(Fig. 3E)**. Together, these observations suggest that cell volume was regulated and that epithelial cells partially compensated geometric constraints so that variations in cell height and apical area offset each other.

As the substrates had very anisotropic curvatures, we next assessed whether cell shapes were anisotropically modulated. Visualization of apical cell outline together with their major and minor axes of elongation revealed that cells adopt anisotropic shapes **(Fig. 3F)**. We then projected the major axis onto the X-Y plane and measured its degree of alignment with the previously defined Y direction. Consistently, we observed a strong curvature-dependent modulation of the alignment: In negative curvature regions (valleys), the major axis of elongation aligned preferentially with the Y axis, whereas in positive curvature regions (crests) cell orientation was markedly more heterogeneous **(Fig. 3G)**.

This orientation bias, together with the curvature-dependent modulation of apical cell area, raised the question of how epithelial cells laterally tile their surfaces as curvature alternates from valley to crest. To address this question, we quantified cell width along the X and Y directions as a function of the periodic coordinate S **(Fig. 3H)**. Strikingly, cell width along the Y axis (valleys/ridge lines) remained largely constant across crests and valleys, and across all substrate geometries. In contrast, cell width along the X axis (valleys to valleys) exhibited pronounced curvature-dependent variations **(Fig. 3I-K)**. These results indicate that cell shape responds to curvature, such that epithelial cells preferentially stretch or compress along the axis of the wavy pattern, while maintaining their dimension along the orthogonal direction.

Collectively, these findings demonstrate that curvature is a powerful geometric cue that remodels epithelial architecture with local curvature primarily driving changes in apical area, anisotropic cell deformation, and cell axis reorientation mostly independently from large-scale curvature. These multiscale responses highlight how epithelia actively remodel to accommodate curved geometries, establishing tissue bending as a key physical regulator of epithelial form.

### Nuclei elongation and biaxiality are conserved across multiscale curvature

We next investigated how multiscale curvature affects nuclear geometry at the three-dimensional level. Confocal imaging of nuclei **(Fig. 4A)** combined with 3D nuclear segmentations **(Fig. 4B)** enabled a systematic quantification of nuclear shape across wavy, parallel and perpendicular substrates. From these segmentations, we first extracted nuclear volume and then performed an ellipsoidal fit giving three main axes of elongation (major, minor 1 and minor 2) and associated radii from the largest to smallest. We computed elongation defined as the aspect ratio (major radius divided by the mean of the minor axes) obtained from ellipsoidal fits **(Fig. 4C top)**, and nuclear biaxiality, quantified as the ratio between the minor 1 and minor 2 ellipsoidal axis **(Fig. 4C bottom)**.

**Figure 4.**
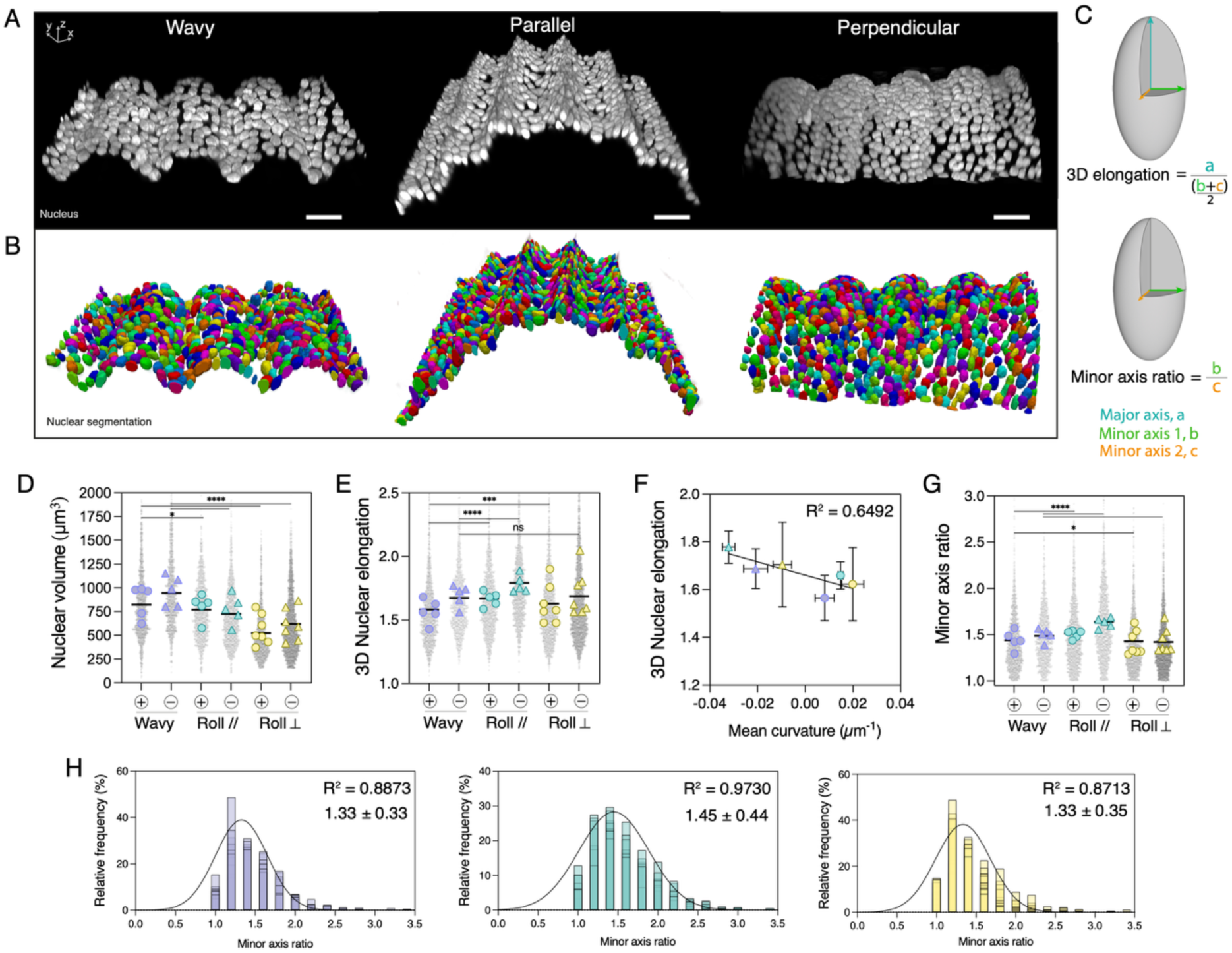
Local curvature modulates nuclear shape and three-dimensional organization. **(A)** Representative confocal images of nuclei in grey and **(B)** the respective nuclear segmentation for wavy, parallel and perpendicular tissues. **(C)** Schematics representation showing the 3D nuclear elongation (top) and the nuclear minor axis ratio (bottom). Quantification of **(D)** the nuclear volume, **(E)** the 3D nuclear elongation and **(G)** the minor axis ratio for positive and negative curvatures of wavy, parallel and perpendicular substrates. **(F)** 3D nuclear elongation is linearly related with the Mean curvature. **(H)** Histogram showing the distribution of minor axis ratio for wavy, parallel and perpendicular substrates. *p<0.05, ***p<0.001 and ****p<0.0001.

We first assessed whether nuclear volume varied in response to curvature modulation **(Fig. 4D)**. Although nuclear volume was slightly reduced in rolled substrates compared to wavy controls, this decrease effect was modest and markedly smaller than the reduction observed in cell area **(Fig. 3B)**, suggesting that nuclear volume is comparatively constrained under multiscale curvature.

To further characterize nuclear shapes, we quantified 3D nuclear elongation **(Fig. 4E)**. Across all substrates, rolling induces only minor changes in nuclear elongation, exhibiting slightly higher aspect ratios in negative curvature (valleys) compared with positive curvature (crests). This trend became emphasized when nuclear elongation was represented as a function of mean curvature in crests and valleys **(Fig. 4F)**. Specifically, nuclear aspect ratios decreased from negative (valleys) to positive (crests) curvature, with value ranging from 1.78 to 1.66 for parallel substrates, 1.71 to 1.62 for perpendicular substrates, and 1.69 to 1.57 for wavy substrates. Despite this curvature-dependent trend, overall nuclear elongation remained generally conserved across conditions.

We next examined nuclear biaxiality by analyzing the ratio between the minor 1 and minor 2 axes of the fitted ellipsoids **(Fig. 4G)**. Strikingly, this ratio remained consistently centered around ∼1.5 across crests and valleys, and across all substrate geometries. As this ratio is by definition larger than one, such a conserved value could in principle arise from fluctuations around a uniaxial nuclear shape. In that case, the most likely value of the ratio should be 1. However, distributions of the minor axis ratio revealed well-defined peaks near ∼1.5 rather than values close to the unity **(Fig. 4H)**, indicating that nuclei preferentially adopt a biaxial, rather than uniaxial, configuration.

While this biaxiality was largely conserved, a modest increase in the minor axis ratio was observed in valleys on parallel substrates, which also correspond to regions of higher negative mean curvature **(Supplementary Fig. 1)**. This suggests that elevated negative curvature can further enhance nuclear biaxiality, albeit within a tightly constrained range.

Collectively, these results demonstrate that nuclei in epithelial monolayers exposed to multiscale curvature adopt a robust and conserved 3D geometry. Nuclei are not only consistently elongated, but also consistently biaxial, with an overall “almond-like” shape in which the major radius exceeds the minor 1 radius, which in turn exceeds the minor 2 radius. This constrained nuclear geometry persists across substrates and curvature conditions, indicating that epithelial nuclei maintain a stereotyped three-dimensional architecture.

### Curvature modulates nuclear alignment and tilt

Having established that nuclear shape is robustly biaxial across substrates, we next asked how the orientation of the nuclear axes vary between negative and positive curvatures. To address this, we developed a dedicated analysis pipeline in which nuclei from different substrates were pooled and grouped into bins (of approximately 8 µm in size) based on their periodic coordinate S **(Supplementary Section 4)**. Nuclear orientation was then expressed in the local reference frame (X,Y,N) defined previously **(Fig. 2C)**, allowing direct comparison of nuclear alignment across positions and substrate geometries.

For each S bin, we constructed an average representative nucleus by averaging the fitted ellipsoids of all nuclei within the bin. The resulting reconstructed average nucleus over one substrate period is shown to scale in **Fig. 5A**. Although individual nuclei displayed substantial orientational variability, this averaging procedure revealed a clear and robust organization. Across all substrate types, the major nuclear axis predominantly lays within the X-Z plane. In valleys, nuclei oriented vertically, with the major axis along the Z direction and the minor 2 axis aligned along the X direction. In contrast, on crests, the behavior of the major axis differed depending on the curvature configuration: in wavy and perpendicular substrates, the major axis rotated from valleys to become nearly horizontal, whereas in parallel substrates it remained largely vertical across the entire substrate.

**Figure 5.**
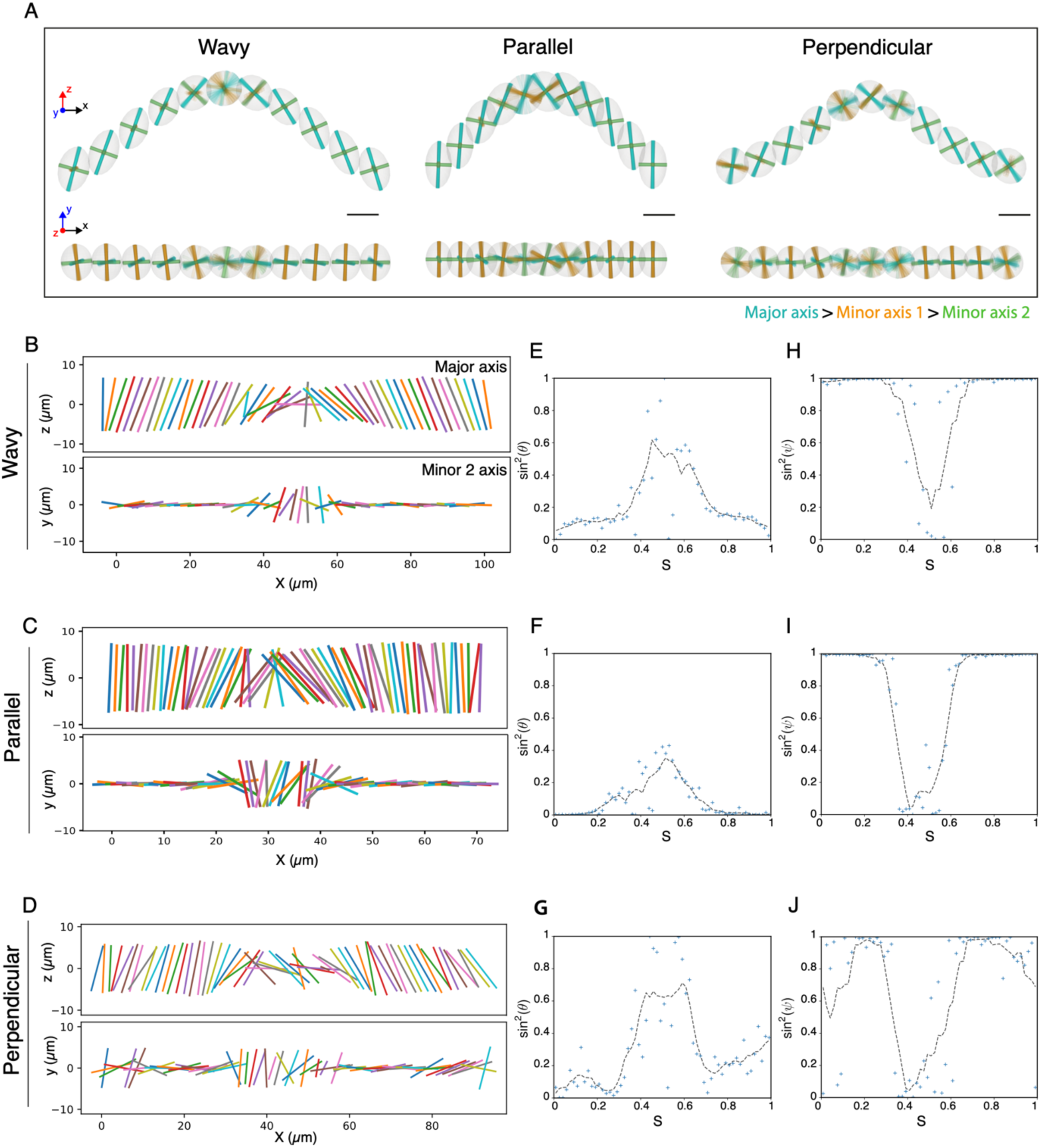
Local curvature directs nuclear orientation within epithelial tissues. **(A)** Average nuclei ellipsoids (grey) of several substrates in the wavy and rolled (parallel, perpendicular) configurations (N=4 substrates for each condition). (Top) X-Z view (Bottom) X-Y view. The major (blue), minor 1 (orange), and minor 2 (green) axes of elongation are shown as solid cylinders, with a blur representing the uncertainty on the estimated average directions. **(B – D)** Representation of the major (Top panels) and minor 2 (Bottom panels) axes orientations in the X-Z (Top panels) and X-Y (Bottom panels) plane respectively, for the wavy **(B)**, parallel **(C)** and perpendicular **(D)** substrates. **(E – G)** Alignment of the major axis with the X direction, quantified by sin^2^ 𝜃, with 𝜃 the angle between the Z axis and the projected major axis. **(H – J)** Alignment of the minor 2 axis with the X direction, quantified by sin^2^ 𝜓, with 𝜓 the angle between the Y axis and the projected minor 2 axis. Blue crosses: Individual bin averages. Dashed lines: Smoothing curves (moving average of 7 nearest points).

To further quantify these orientation patterns, we analyzed nuclear axis alignment at finer spatial resolution along *S*. We first considered the orientation of the major axis projected onto the X-Z plane and the minor axis projected onto the X-Y plane as a function of X **(Fig. 5B-D)**. We define the angle 𝜃 as the angle between the projected major axis and the Z axis. Plotting sin^2^ 𝜃 as a function of S revealed **(Fig. 5E-G)** that, for wavy **(Fig. 5E)** and perpendicular **(Fig. 5F)** substrates, the major axis became fully horizontal at S=0.5, corresponding to crest (positive curvature) regions. In contrast, in parallel substrates **(Fig. 5F)**, the major axis exhibited bending but did not fully flatten, remaining partially aligned with the vertical direction.

We next quantified the orientation of the minor axis by projecting it onto the *X–Y* plane and defining the angle 𝜓 between the projected minor axis and the Y axis. Across all substrates, sin^2^ 𝜓 reached its minimal value, close to 0, at S ≈ 0.5 **(Fig. 5H-J)**, indicating that the minor axis consistently reoriented toward the Y direction around the crests.

Together, these findings demonstrated that nuclear orientation is strongly modulated by curvature in a position- and geometry-dependent manner. Nuclear alignment emerges from the interplay between local geometric constraints and collective packing, with curvature favoring alignment along principal directions of curvature while neighboring nuclei impose steric constraints that promote orientational coherence. Within this line of reasoning, the persistence of vertically oriented nuclei the parallel substrates may arise from the reduced oscillation wavelength in this configuration **(Fig. 1H)**, which likely increases the energetic cost associated with bending or reorienting elongated nuclei.

## Discussion

In this study, we establish a microfabricated platform that combines wavy hydrogels with inducible self-rolling substrates to independently impose local and large-scale curvature on epithelial monolayers. Using this platform, we simultaneously impose different local and large-scale curvatures on epithelial monolayers and study the regulation of epithelial organization across multiple curvature scales. We imposed well-defined curvature patterns on epithelial monolayers while preserving homogeneous material properties and biochemical conditions, allowing curvature-specific effects to be isolated. Using this system, we reveal how tissue geometry alone can drive coordinated changes in cell architecture, from cell packing to nuclear organization.

Consistent with previous observations, curvature induction via the self-rolling system imposes a certain degree of compression during rolling^24^. In the presence of a wavy hydrogel layer, however, this compression is redistributed in a more complex, orientation-dependent manner. Indeed, the wavy surface increases resistance to self-rolling, resulting in larger tube radii overall, and particularly in perpendicular substrates, indicating reduced effective compression.

At the tissue and cellular levels, our results demonstrate that curvature imposes different morphological changes and directional mechanical constraints that reorganize epithelial packing. On negative curvature, cells exhibit a reduction in apical cell area and a pronounced thickening whereas on positive curvature, cells appear wider and less thick in agreement with previous observations^9,12,28^. This suggests an anti-correlation between apical cell area and tissue height which is consistent with a regulation of cell volume at the tissue level and responding to the local and large-scale curvature.

Strikingly, we also demonstrated that MDCK cells adopt anisotropic shapes along the X and Y axes, which was not previously shown^9^. The cell major axis aligned perfectly with the Y axis in negatively curved regions, suggesting that the cell elongation and orientation respond strongly, and only, to local curvature. This anisotropic response indicates that curvature generates spatially heterogeneous in-plane stresses that bias cell shape remodeling. As previously demonstrated, on positively curved regions, HK-2 cells aligned their in-plane axis perpendicular to the channel. However, on negatively curved regions, cells were elongated and showed longitudinal directionality^29^. Such behavior aligns with the fact that curved surfaces redistribute cortical tension and junctional forces to guide collective cell organization.

Beyond cell shape, our data reveal that curvature-dependent mechanical cues propagate to the nucleus. As demonstrated before, the nucleus can be considered as a topographical sensor and a mechanical guide^30,31^. In this study, we showed first that nuclear volume remains relatively stable suggesting that curvature primarily affects nuclear shape rather than inducing compressive volume loss. Indeed, we found that nuclei preferentially adopt an elongated state on all types of substrates with a slight increase on negative curvature. In addition to the 3D nuclear elongation, we demonstrate that the minor axis ratio of nuclei is always around 1.5 for each substrate meaning that nuclei have a biaxial configuration. Strikingly, we found that the main effect of curvature is focused on nuclei orientation, instead of nuclei morphologies. Nuclei reorient in response to local curvature, indicating a high sensitivity to tissue-scale geometric constraints. Given that nuclear deformation can trigger changes in cell state^32^ through modulations of chromatin architecture and transcription^33^, these orientation changes raise the possibility that curvature may indirectly pattern gene expression through nuclear alignment. In physiological contexts such as tubular organs, intestinal folds, or developing epithelia, spatial variations in curvature could therefore encode patterned mechanical signals that contribute to regionalized cellular states.

Finally, the dual-scale curved substrates presented here provide a versatile framework to investigate curvature-driven mechanobiology. Unlike traditional micropatterned or stretch-based systems, this platform enables complex curvature landscapes with controlled orientation relative to tissue axes, closely mimicking in vivo geometries. This versatility opens opportunities to explore how curvature interacts with other mechanical parameters, such as substrate stiffness, contractility, or tissue thickness, and how curvature sensing evolves over time during growth, remodeling, or disease progression.

## Conclusion

In conclusion, our findings identify curvature as a fundamental geometric regulator of epithelial organization, linking tissue-scale architecture to cellular and nuclear organization. By combining experimental and computational approaches, we show that epithelial architecture emerges from local mechanical interactions in which curvature governs the balance between apical expansion, volume conservation, and tissue thickness. These findings suggest that large-scale tissue morphogenesis can arise from generic physical constraints at the cellular level, without requiring additional biochemical regulation of cell shape. Future studies integrating molecular readouts and dynamic perturbations will be essential to determine how curvature-driven nuclear remodeling translates into functional changes in gene expression and cell fate

## Materials and methods

### Self-rolling wavy substrate preparation

#### Self-rolling substrates

Polydimethylsiloxane (PDMS, Sylgard 184 from Dow Corning) mixture of 10:1 monomer:crosslinker ratio was spin-coated on circular glass coverslips previously washed in a 70% ethanol solution during 15min. PDMS was then cured at 60 °C for 4 hours to obtain a thin membrane of 3 MPa and ∼60 μm thick. This PDMS layer serves to obtain a sharp cut of material above it, allowing the blade to lean on it. The PDMS surface was then activated using a plasma cleaner for 45s, though a 0.5 cm high PDMS mask with a central hole of 0.5cm by 1cm. 15µl of a 3% wt/vol of fish gelatin water solution were incubated 2h on the activated surface. Then, approx. 100µl of a solution of PDMS Sylgard 184, curing agent and toluene mixed in a 10:1:10 proportion were spin-coated onto the gelatin/PDMS surface (600rpm/s, 3000rpm, 50s), and reticulated on a hotplate at 110° for 30min. Then, approx. 100µl of a second solution of PDMS Sylgard 184, curing agent, silicon oil and toluene mixed in a 5:5:0.5:10 proportion were spin-coated (600rpm/s and, 1500 rpm, 50s), and reticulated. The substrate was incubated in isopropanol overnight to extract the silicon oil. Finally, coverslips were dried on a hotplate at 80° for 5min before storage ^24^.

#### Binding PDMS layers and wavy polyacrylamide hydrogel

The PDMS substrates were incubated in a solution of 10 wt%/vol benzophenone (Sigma) dissolved in a water/acetone mixture (35: 65 w/w) during 60s. Substrates were immediately rinsed with methanol during 60s and dried with nitrogen. The benzophenone-infused PDMS substrate was plasma activated for 45s. This step needs to be performed a few minutes before UV illumination to maintain hydrophilicity. Wavy polyacrylamide hydrogel solution was prepared according to previously reported methods ^9^. Briefly, to form a composite network of acrylamide and PDMS, we prepared a solution by mixing acrylamide (AAm), bis-acrylamide (bis-AAm), N-hydroxyethylacrylamide (HEA), 2-Hydroxy-4′-(2-hydroxyethoxy)-2 methylpropiophenone (Irgacure 2959) and deionized water. 20µl of this solution was pipetted onto pattern of chromium optical photomask (Toppan photomask, France) and covered with the PDMS substrate. The system was exposed to UV illumination at 360nm during 8min30 (Dymax UV light curing lamp). In this study, we used transparent stripes of 25 µm wide and black stripes of 75 µm form isotropic wavy hydrogels with wavelengths of 100 µm. Finally, wavy-PDMS substrates were gently removed from the photomask under water immersion, washed three times in sterile deionized water under gentle agitation and stored in sterile deionized water at 4°C.

### Cell culture and samples preparation

Epithelial cells from the Madin-Darby Canine Kidney cell line (MDCK II, Sigma #85011435) were maintained in polystyrene T75 flasks of in a cell culture incubator at 37°C and 5% CO^2^. MDCK cells were cultured in proliferation medium composed of Dubelcco’s Modified Eagle’s medium (DMEM), high glucose (4.5 g/l) with L-glutamine (BE12-604F, Lonza) supplemented with 10% (v/v) Fetal Bovine Serum (FBS, AE Scientific) and 1% of penicillin and streptomycin antibiotics (AE Scientific). The hydrogel surface was incubated with fibronectin (FN, Merck) at 75µg/ml in sterile deionized water, during 1h to allow cell adhesion. MDCK cells were seeded onto the fibronectin-coated hydrogel surface at a density of 2000 cells/mm^2^ for 48 hours.

#### Cells on wavy self-rolling substrates

By crosswise cutting the hydrogel-PDMS system with a scalpel in the central region, the gelatin swells and dissolves upon contact with the warm cell culture medium, releasing the pre- constraint and allowing for PDMS rolling. Thus, 2 symmetric parallel and 2 symmetric perpendicular tubes were produced per sample, kept in the incubator during 5min after tube formation, and then fixed and permeabilized with 4% paraformaldehyde (PFA, Electron Microscopy Sciences) and 0.05% Triton X-100 (Sigma) in phosphate buffered saline (PBS 1X, Capricorn scientific) for 20 min at room temperature, washed three times with PBS and kept in PBS at 4^°^C until imaging.

### Immunostaining of epithelial tissues

Cell membrane was labelled on live MDCK-II cells with CellMask^TM^ Deep Red Plasma Membrane Dye (1:3000) incubated for 30 min in a fresh medium and nuclei were labelled with Hoechst 33342 (1:2000) for 15 min. Cells were then fixed 5 minutes after cutting of the self-rolling substrates. After fixation, coverslip with immunostained cells were immerged in glycerol for confocal imaging.

### Confocal imaging

MDCK epithelial tissues were observed in confocal mode with an upright LSM 710 microscope with a water immersion Plan-Apochromat 20x/1.0 N.A. DIC objective (Carl Zeiss, Oberkochen, Germany), and operated with Zen 2012 software. Z-stack acquisitions were performed with Z-step of 0.5 mm for cell imaging. A Nikon A1R HD25 Ti2 motorized inverted microscope (Nikon, Japan) with x60 and 100x objectives was used to perform more detailed confocal images using small Z-depth increments (0.15 μm) and for imaging of substrates. Confocal images were recorded and processed using NIS-Elements (Nikon, Advanced Research v.4.5).

### Morphometric analysis of the nuclei

#### Nuclei segmentation

Nuclei segmentation was performed using NIS-Element Analysis v.5.42.06 software with General Analysis 3 (GA3). This versatile instrument is designed to construct image analysis procedures in a modular manner. Images with fluorescent nuclei labelled with Hoechst were segmented via a homemade pipeline with GA3 giving the nuclear volume.

#### Matlab script

From the NIS-Element Analysis software, we obtained a 3D array of voxels where every voxel is tagged as being either part of a specific nuclei, or of the background. The volume was simply extracted via the number of voxels per nuclei. The elongations were obtained by first identifying voxels belonging to the nuclei’s surface, based on their connectivity with background voxels. The primary axis of elongation was then extracted by finding the two most distant nuclei surface voxels. The secondary axis of elongation was then defined by finding the nuclei surface voxels furthest away from the primary axis of elongation. Lastly, the third axis of elongation was defined as the vector perpendicular to the first two axis of elongation and passing by the intersection of the first two axis of elongation.

### Surface segmentation

#### Global shape extraction and cell segmentation

The extraction of global shape was performed with MorphographX (MGX) software. MGX is an open-source platform for the visualization and processing of 3D image data. A cell outline marker is necessary to proceed MGX steps. MGX was used to segment both apical and basal surface. Briefly, volumetric data is pre-processed to remove noise from surface. The next step is to define a surface following the global shape of the tissue with the edge detect function leading to have a mask (binary image) of either apical or basal surface. The surface is extracted as an apical or basal surface mesh. After defining this mesh, the membrane signal just below the surface mesh (between 1 and 3µm away) can be projected onto the surface mesh, creating a curved image of the outer layer of cells. The signal is then segmentate via a watershed segmentation. This enables the extraction of precise cell outlines without the distortions associated with a flat 2D projection. Cell area and cell outlines can be extracted from the software ^34^.

### Statistical analysis

The statistical significance of differences between conditions was analyzed using Prism v.10 (GraphPad Software, Inc). For multiple comparisons the differences were determined by using an analysis of variance, a Kruskal-Wallis test. Differences with a p-value under 0.05 were considered statistically significant. Significance is indicated by asterisks in figures (*p<0.05, **p<0.01, ***p<0.001, ****p<0.0001) and n.s. means not significant. Unless otherwise stated, all data are presented as mean ± standard deviation (S.D.).

## Acknowledgments

A.R. acknowledges funding from the Swiss National Fund for Research grant numbers #CRSII5_189996 and #310030_200793 and the European Research Council Synergy grant number #951324-R2-TENSION. C.T. acknowledges support from the French National Research Agency, grant no. ANR-22-CE13-0015-01 for the project “CurvEDyn”. M.L. is Postdoctoral Fellow (Chargée de Recherches) of the National Fund for Scientific Research (F.R.S-FNRS).

## Author contributions

**(A)** M. L., Q.V., G.S., C.T. and A.R. designed the research. M.L. and C.T. established the wavy rolling system. M.L. performed all of the experiments and image analyses. G.P. developed a homemade Matlab code for nuclei properties extraction. Q.V., with inputs of G.P., constructed the analysis pipeline. Figure designs were generated by all authors and further edited by M.L and Q.V. M.L. and Q.V. analyzed the data. M.L., Q.V., G.S. and A.R. wrote the paper with contributions from all authors.

---

**Supplementary Figure 1.**
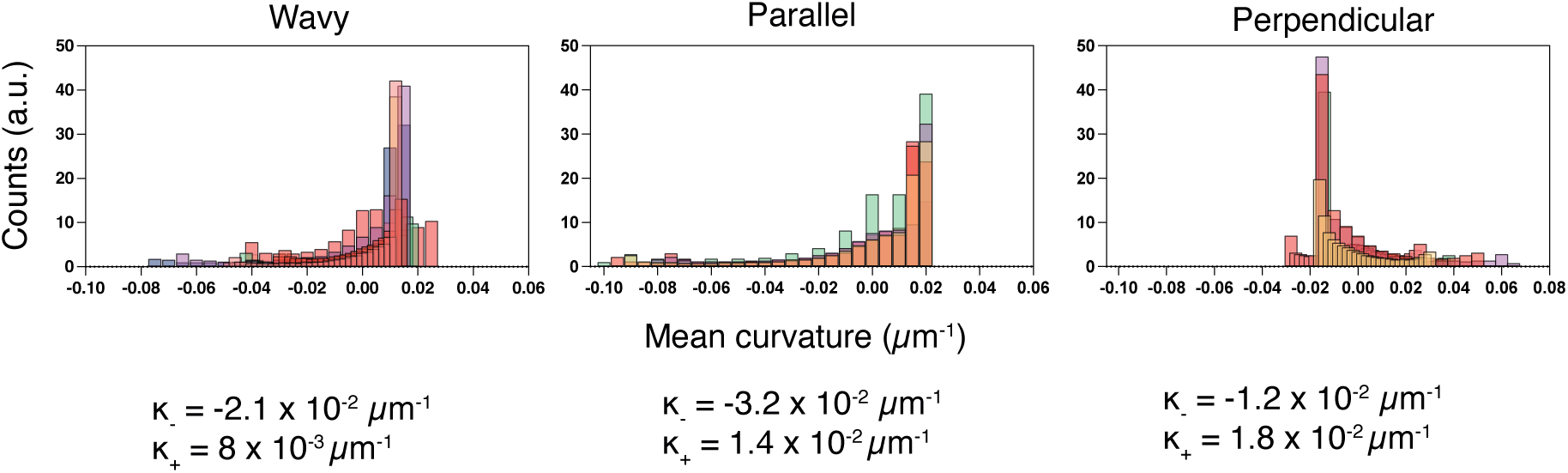
Distribution of mean curvature across wavy and rolled substrates. Histograms of mean curvature for wavy, parallel and perpendicular structures (N = 7, 5, 5 substrates for wavy, parallel and perpendicular). 𝜅_–_ and 𝜅_+_ correspond to the mean of negative and positive curvature, respectively.

**Supplementary Figure 2.**
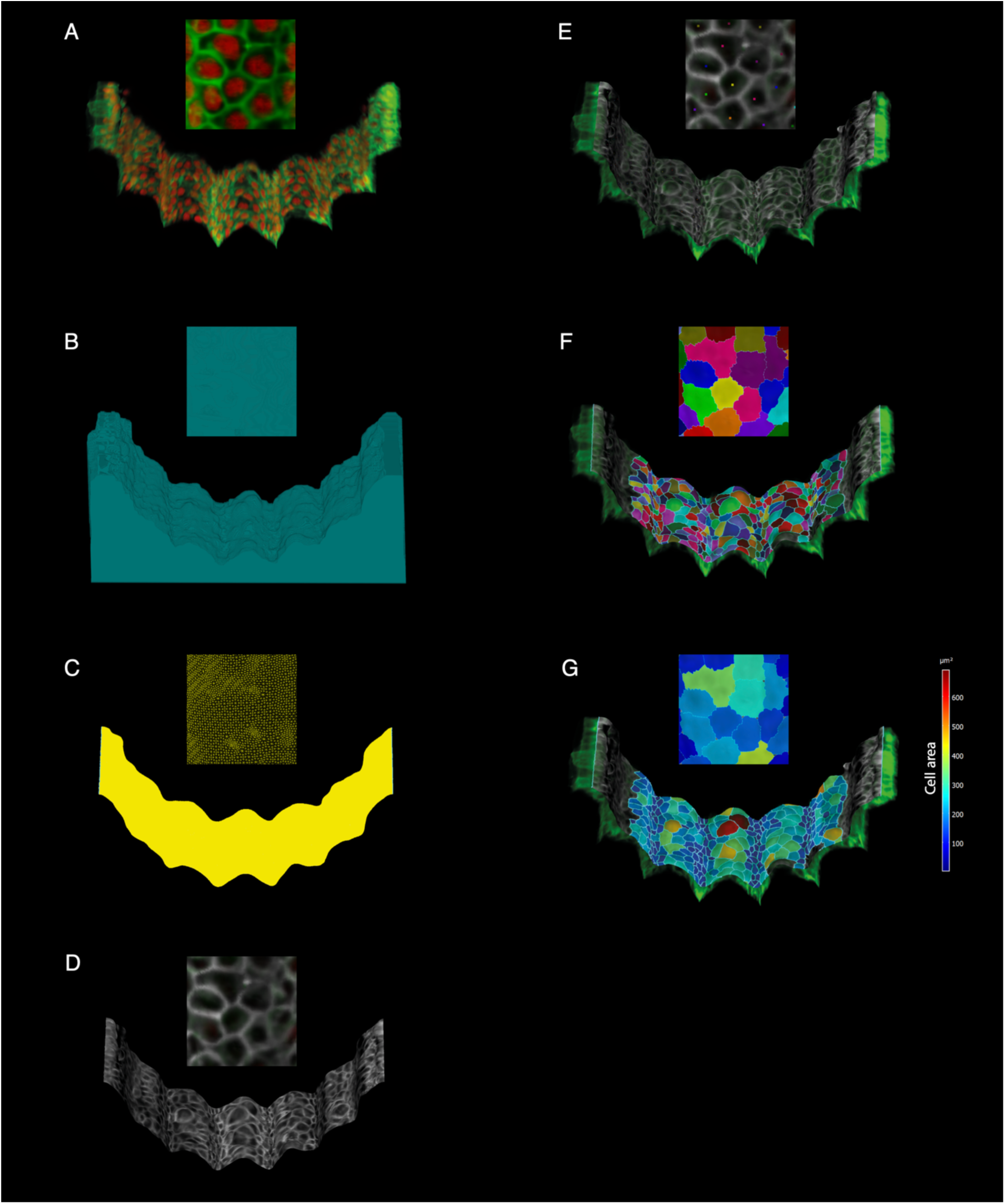
Automated 3D morphometric pipeline for curvature analysis using MorphoGraphX. (A) Raw 3D fluorescence of cell membrane (green) and nuclei (red). **(B)** Extraction of global shape of the curved monolayer. **(C)** The surface of the object is then extracted from this shape as a triangular mesh to create a mesh (in yellow). **(D)** Projection of the fluorescent signal onto the mesh. **(E-F)** Cell segmentation is done by propagating label seeds on the mesh surface using the watershed algorithm. Label seeds are placed automatically. **(G)** Heat-map of cell area across rolled surface. Scale bar, 100µm.

## References

1. Arnold, C. et al. Bending the rules: curvature’s impact on cell biology. BMC Biol 23, 296 (2025).

2. Schamberger, B. et al. Curvature in Biological Systems: Its Quantification, Emergence, and Implications across the Scales. Advanced Materials 35, 2206110 (2023).

3. Wang, Y., Stonehouse-Smith, D., Cobourne, M. T., Green, J. B. A. & Seppala, M. Cellular mechanisms of reverse epithelial curvature in tissue morphogenesis. Front. Cell Dev. Biol. 10, (2022).

4. Khoromskaia, D. & Salbreux, G. Active morphogenesis of patterned epithelial shells. eLife 12, e75878 (2023).

5. Luciano, M., Tomba, C., Roux, A. & Gabriele, S. How multiscale curvature couples forces to cellular functions. Nat Rev Phys 6, 246–268 (2024).

6. Tanabe, N., Sato, S., Suki, B. & Hirai, T. Fractal Analysis of Lung Structure in Chronic Obstructive Pulmonary Disease. Front Physiol 11, 603197 (2020).

7. Belchi, F. et al. Lung Topology Characteristics in patients with Chronic Obstructive Pulmonary Disease. Sci Rep 8, 5341 (2018).

8. Luciano, M. et al. Appreciating the role of cell shape changes in the mechanobiology of epithelial tissues. Biophysics Rev. 3, 011305 (2022).

9. Luciano, M. et al. Cell monolayers sense curvature by exploiting active mechanics and nuclear mechanoadaptation. Nat. Phys. 17, 1382–1390 (2021).

10. Luciano, M., Versaevel, M., Kalukula, Y. & Gabriele, S. Mechanoresponse of Curved Epithelial Monolayers Lining Bowl-Shaped 3D Microwells. Adv Healthcare Materials 13, 2203377 (2024).

11. Huang, C.-K., Yong, X., She, D. T. & Lim, C. T. Surface curvature and basal hydraulic stress induce spatial bias in cell extrusion. eLife 12, (2024).

12. Werner, M. et al. Surface Curvature Differentially Regulates Stem Cell Migration and Differentiation via Altered Attachment Morphology and Nuclear Deformation. Adv. Sci. 4, 1600347 (2017).

13. Leclech, C. et al. Micro-Scale Topography Triggers Dynamic 3D Nuclear Deformations. Advanced Science 12, 2410052 (2025).

14. Pieuchot, L. et al. Curvotaxis directs cell migration through cell-scale curvature landscapes. Nat Commun 9, 3995 (2018).

15. Ravasio, A. et al. Gap geometry dictates epithelial closure efficiency. Nat Commun 6, 7683 (2015).

16. Senger, F. et al. Spatial integration of mechanical forces by α-actinin establishes actin network symmetry. Journal of Cell Science 132, jcs236604 (2019).

17. Chen, T. et al. Large-scale curvature sensing by directional actin flow drives cellular migration mode switching. Nat. Phys. 15, 393–402 (2019).

18. Luciano, M. & Gabriele, S. Designing hydrogel dimensionality to investigate mechanobiology. Soft Matter 21, 4551–4572 (2025).

19. Pahapale, G. J. et al. Directing Multicellular Organization by Varying the Aspect Ratio of Soft Hydrogel Microwells. Advanced Science 9, 2104649 (2022).

20. Glentis, A. et al. The emergence of spontaneous coordinated epithelial rotation on cylindrical curved surfaces. Sci. Adv. 8, eabn5406 (2022).

21. Bril, M. et al. Dynamic substrate topographies drive actin- and vimentin-mediated nuclear mechanoprotection events in human fibroblasts. BMC Biol 23, 94 (2025).

22. Ouedraogo, S. et al. Fabrication and characterization of thin self-rolling film for anti-inflammatory drug delivery. Colloids and Surfaces B: Biointerfaces 241, 114039 (2024).

23. Egunov, A. I., Korvink, J. G. & Luchnikov, V. A. Polydimethylsiloxane bilayer films with an embedded spontaneous curvature. Soft Matter 12, 45–52 (2015).

24. Tomba, C., Luchnikov, V., Barberi, L., Blanch-Mercader, C. & Roux, A. Epithelial cells adapt to curvature induction via transient active osmotic swelling. Developmental Cell 57, 1257–1270.e5 (2022).

25. Kalukula, Y. et al. Unlocking the therapeutic potential of cellular mechanobiology. Science Advances 11, eaea6817 (2025).

26. Messal, H. A. et al. Tissue curvature and apicobasal mechanical tension imbalance instruct cancer morphogenesis. Nature 566, 126–130 (2019).

27. Simmons, C. S., Ribeiro, A. J. S. & Pruitt, B. L. Formation of composite polyacrylamide and silicone substrates for independent control of stiffness and strain. Lab Chip 13, 646 (2013).

28. Ehrig, S. et al. Surface tension determines tissue shape and growth kinetics. Sci Adv 5, 7 (2019).

29. Yu, S.-M. Substrate curvature affects the shape, orientation, and polarization of renal epithelial cells. Acta Biomaterialia 77, 311–321 (2018).

30. Vassaux, M., Pieuchot, L., Anselme, K., Bigerelle, M. & Milan, J.-L. A Biophysical Model for Curvature-Guided Cell Migration. Biophys J 117, 1136–1144 (2019).

31. Niethammer, P. Components and Mechanisms of Nuclear Mechanotransduction. Annu Rev Cell Dev Biol 37, 233–256 (2021).

32. Versaevel, M., Grevesse, T. & Gabriele, S. Spatial coordination between cell and nuclear shape within micropatterned endothelial cells. Nat Commun 3, 671 (2012).

33. Kalukula, Y., Stephens, A. D., Lammerding, J. & Gabriele, S. Mechanics and functional consequences of nuclear deformations. Nat Rev Mol Cell Biol 23, 583–602 (2022).

34. Barbier de Reuille, P., et al. MorphoGraphX: A platform for quantifying morphogenesis in 4D. eLife 4, e05864 (2015).

